# ShiftSCAN, a program that predicts potential alternative sources of mass spectrometry-derived peptides, improves the accuracy of studies on novel amino acid sequences

**DOI:** 10.1101/2025.05.30.656965

**Authors:** Umut Çakır, Noujoud Gabed, Ali Yurtseven, Igor Kryvoruchko

## Abstract

Mass spectrometry (MS) proteomics is currently the most powerful tool for identifying both annotated proteins and proteins translated from non-canonical open reading frames or unusual genetic events. With this method, numerous novel protein-coding loci have been discovered by searching for short fragments of hypothetical longer peptides and polypeptides. Apart from the validation of translation from mRNA transcripts, MS proteomics has been instrumental for the detection of peptides encoded by non-mRNA transcripts. A special application field of MS proteomics is studies on programmed ribosomal frameshifting (PRF), where the detection of chimeric peptides produced from two different reading frames is vital. Each novel chimeric peptide is thought to originate from a certain genetic locus. However, due to the short length of MS peptides, there is a possibility that MS-supported chimeric peptides are produced by additional (alternative) loci via PRF. This scenario evaded due attention because the contribution of non-canonical peptides and proteins to the functional diversity of proteomes is still thought to be minor. Recent studies have challenged this paradigm. To the best of our knowledge, our group was the first to include alternative chimeric sources into the analysis pipeline. This resulted in a much higher certainty about loci responsible for the production of unusual peptides. This certainty is crucial for the functional characterization of such loci. At the same time, our study revealed enormous diversity of potential alternative sources for a subset of MS-supported non-canonical peptides. Here, we present a highly flexible program that predicts alternative chimeric and non-chimeric sources of peptides detected by MS proteomics.

## 1. Introduction

Translation of peptides and proteins that deviate from expected products of annotated, or reference, open reading frames (refORFs) is defined as non-canonical (Wright et al., 2022). Large-scale detection of non-canonical translation events is important for understanding the full complexity of proteomes (Landry et al., 2015). Translated alternative open reading frames (altORFs) and their products, alternative proteins (altProts), have been recognized as a major source of protein diversity (Mouilleron et al., 2016). Many of them have been shown to have vital functions (Orr et al., 2020). A dedicated database of altORFs has been created for several species including humans (Brunet et al., 2019, 2021; Leblanc et al., 2024). Mass spectrometry (MS) proteomics in conjunction with ribosome profiling and conservation analysis serves as a major tool for the detection of biologically relevant altORFs (Wright et al., 2022). AltORFs can be translated from mRNA transcripts, where they may overlap with the annotated coding sequences (Sheshukova et al., 2017). They can also be translated from other types of RNA (Orr et al., 2020). In this category, long non-coding RNA (lncRNA) is the best-characterized source of translated altORFs (Ruiz-Orera et al., 2014, 2020; Ji et al., 2015; Bazin et al., 2017; Sruthi et al., 2022). Because altORFs are typically shorter than 300 nt, they are often called short ORFs (sORFs) or upstream ORFs (uORFs), if they are located in the 5’-untranslated regions (Orr et al., 2020). Historically, the ORF length alone was the key criterion in genome annotation pipelines. For that reason, the longest ORF found in a transcript is usually considered as a refORF, which sometimes is very misleading especially because altORFs often bear striking similarity to their refORFs (Mouilleron et al., 2016; Wright et al., 2022; Wang et al., 2022). There are still very few translated altORFs known in transcript types other than mRNA and ncRNA. Documented examples of rRNA (Maximov et al., 2002), pre-miRNA (Wang et al., 2014; Couzigou et al., 2016), and pre-siRNA transcripts (Yoshikawa et al., 2016) acting as templates for ribosomes are available in the literature. In addition to translation to discrete altProts, altORFs that overlap with refORFs or other altORFs can be translated with the involvement of programmed ribosomal frameshifting (PRF). Products of such translation are called chimeric peptides and proteins to discriminate them from other types of altProts (Ketteler, 2012; Riegger and Caliskan, 2022; Çakır et al., 2023). The reports about chimeric peptides found in non-viral systems have been known since 1985 (Craigen et al., 1985; Craigen and Caskey, 1986). However, relatively few genetic loci that use PRF for translation in prokaryotes (Blinkowa and Walker, 1990; Flower and McHenry, 1990; Tsuchihashi and Kornberg, 1990; Chaijarasphong et al, 2016; Meydan et al., 2017) and eukaryotes (Matsufuji et al., 1995; Clark et al., 2007; Ivanov and Atkins, 2007; Ren et al., 2024) have been studied so far. Viruses have to use their limited genomic spaces very efficiently by involving all six reading frames and PRF in their translation. Non-viral genomes, in contrast, have no major constraint on sizes. Thus, PRF is often thought to play no major role in cellular processes. Earlier bioinformatic predictions (Ketteler, 2012) and two recent reports challenged this paradigm. A study in humans found 405 unique MS-supported chimeric peptides in 32 samples under normal pathophysiological conditions (Ren et al., 2024). Our computational study in the model plant *Medicago truncatula* identified 156 putative chimeric peptides in three MS proteomic datasets from 16 normal and stressed biological samples (Çakır et al., 2024, preprint). In addition to chimeric peptides produced from mRNA and lncRNA, we also found evidence for chimeric translation from rRNA and tRNA. Because the large-scale production of chimeric peptides is not expected under the dominating assumption of no importance of PRF beyond viruses, it is crucial to trace the origin of each detected chimeric MS peptide back to specific loci where putative PRF sites can be located. Without the knowledge about all loci that can potentially produce the same chimeric sequence, possibly by different kinds of PRF events (Figure 1), there is intrinsic ambiguity that precludes any genetic studies on such loci. Namely, if a given MS-supported chimeric peptide can theoretically come from multiple chimeric sources, the relative contribution of such sources to the translation of that chimeric peptide must be assessed thoroughly. This assessment can be based on the accordance between transcriptome and proteome samples, and also on the relative magnitude of expression in cells, cell types, tissues, organs, and/or conditions relevant to the MS-based detection (Çakır et al., 2024, preprint). In other words, when a novel chimeric peptide is identified via MS proteomics, the key questions are whether this peptide can be produced (1) without PRF from any location in the genome and/or (2) via PRF from any location other than the primary-source locus (the locus from which it has been discovered). The first question is easily addressed by a standard TBLASTN algorithm applied to nucleotide sequences (the annotated transcriptome, the RNA-Seq-derived *de facto* transcriptome, and the repeatome or the transposome). In contrast, no tool has been available so far for the identification of alternative chimeric sources of putative chimeric peptides. Importantly, such a tool can also be very useful for resolving ambiguity cases of the opposite type: a proof that a putative non-chimeric sequence (i.e., no frameshift involved) cannot be produced by a PRF event. This unusual situation can be faced when a novel translated altORF is detected in a transcript in which it is highly unexpected, for example, any transcript that does not fall into the category of mRNA and lncRNA. Our script presented here was designed specifically to address these scenarios. Here we describe the successful application of our software for the analysis of 156 MS-supported chimeric peptides in *M. truncatula*. Although large-scale detection of chimeric peptides is still very rare, we hope that our tool will be of high value for such studies in the future.

**Figure 1.**
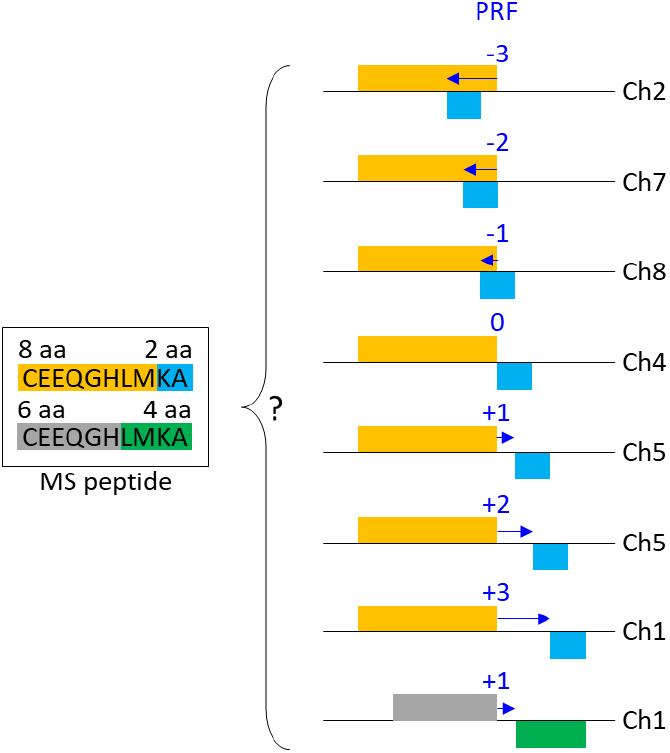
This graph illustrates the ambiguity about the true genetic origin of a mass-spectrometry (MS)-detected chimeric peptide. In this example, a 10-aa-long MS peptide was discovered as a product of a programmed ribosomal frameshifting (PRF) event with value -2 at a locus on Chromosome 7 (Ch7). The PRF event divides the peptide into two arms. The left arm shown in orange contains 8 aa and the right arm (blue) contains 2 aa. However, the same peptide could potentially be produced via a -3 frameshift on Chromosome 2, a -1 PRF event on Chromosome 8, etc., and even without any frameshift (PRF value 0) on Chromosome 4. At the same time, a completely identical amino acid sequence may correspond to a +1 PRF event on Chromosome 8 that divides the peptide into two different arms (gray and green). ShiftSCAN identifies all alternative chimeric and non-chimeric sources of MS peptides, taking into consideration hypothetical PRF events with any value from 0 to infinity. It also identifies potential sources of MS peptides on a complementary strand of a nucleotide sequence, for example, on a reverse complement of a transposon.

## 2. Material and methods

ShiftSCAN stands for Shift detection and Source Confirmation by Alignment and Navigation. It can predict alternative chimeric and non-chimeric sources of translation using two input files: subject database and query database. The subject database can contain any deoxyribonucleotide sequence regardless of its type, origin, and translation status. The query database can be composed of chimeric or non-chimeric amino acid sequences, depending on the purpose of the analysis. For each peptide, the algorithm virtually divides the sequence into two segments: a left segment and a right segment. First, ShiftSCAN searches for both segments in the subject sequence. If both are identified within the distance defined by the setting “gap” (2 in our analysis), the distance is converted into a frameshift (PRF) value. For example, if the gap is 2 (two segments are spaced by 2 nt), the PRF value is reported as +2. If the gap is minus 2 nt (two segments overlap by 2 nt), the reported PRF value is -2. The length of the left segment can range between one amino acid and the full length of the query sequence minus one amino acid. But it can also be the full query sequence, if the alternative source involves no frameshift. Critically, the distance between the left and right matches can vary between zero and a user defined value, simulating possible non-canonical ribosomal slippage events. A gap of zero nucleotides corresponds to uninterrupted translation (non-chimeric sources). ShiftSCAN records detailed information about the event, including the presence or absence of a frameshift, the PRF value, the PRF type (e.g., frame 1 → frame 2), the position of the frameshift within the peptide, the title and sequence of the matched nucleotide subject, and the reading frame direction. All output parameters generated by the program are described in detail in Table 1.

**Table 1.**
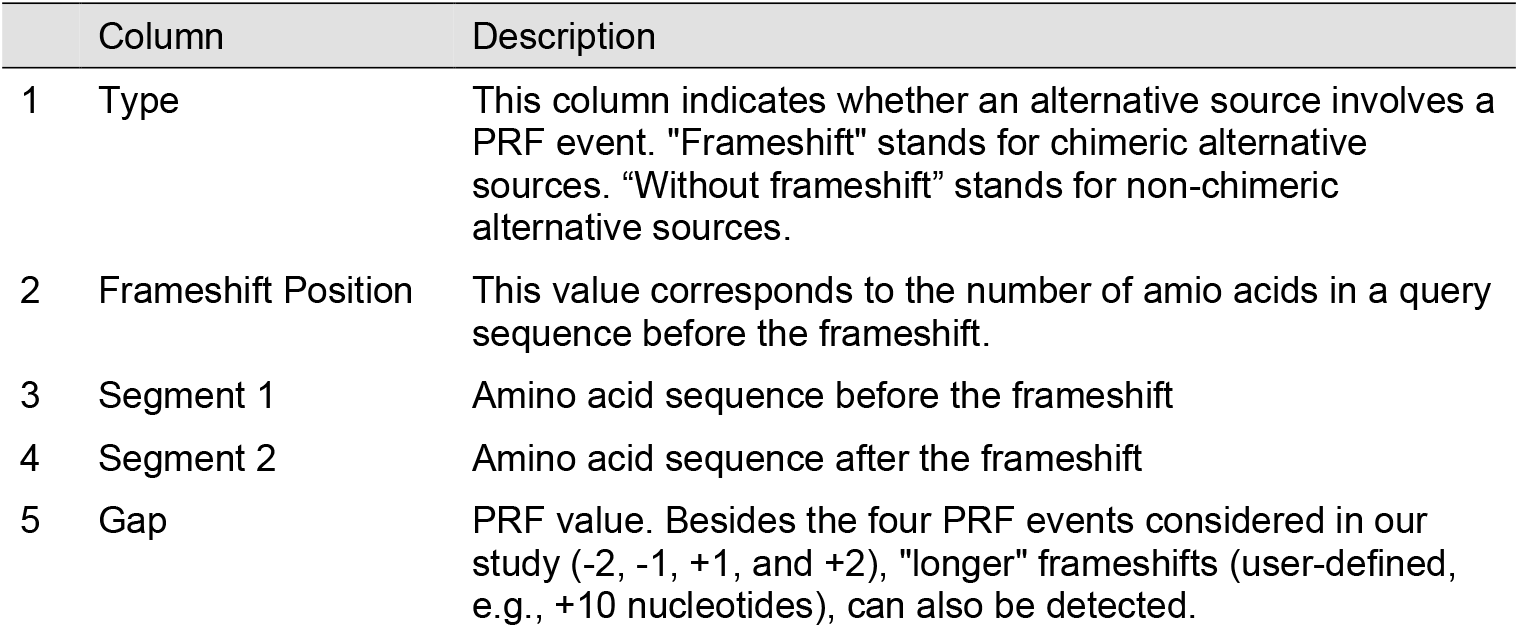
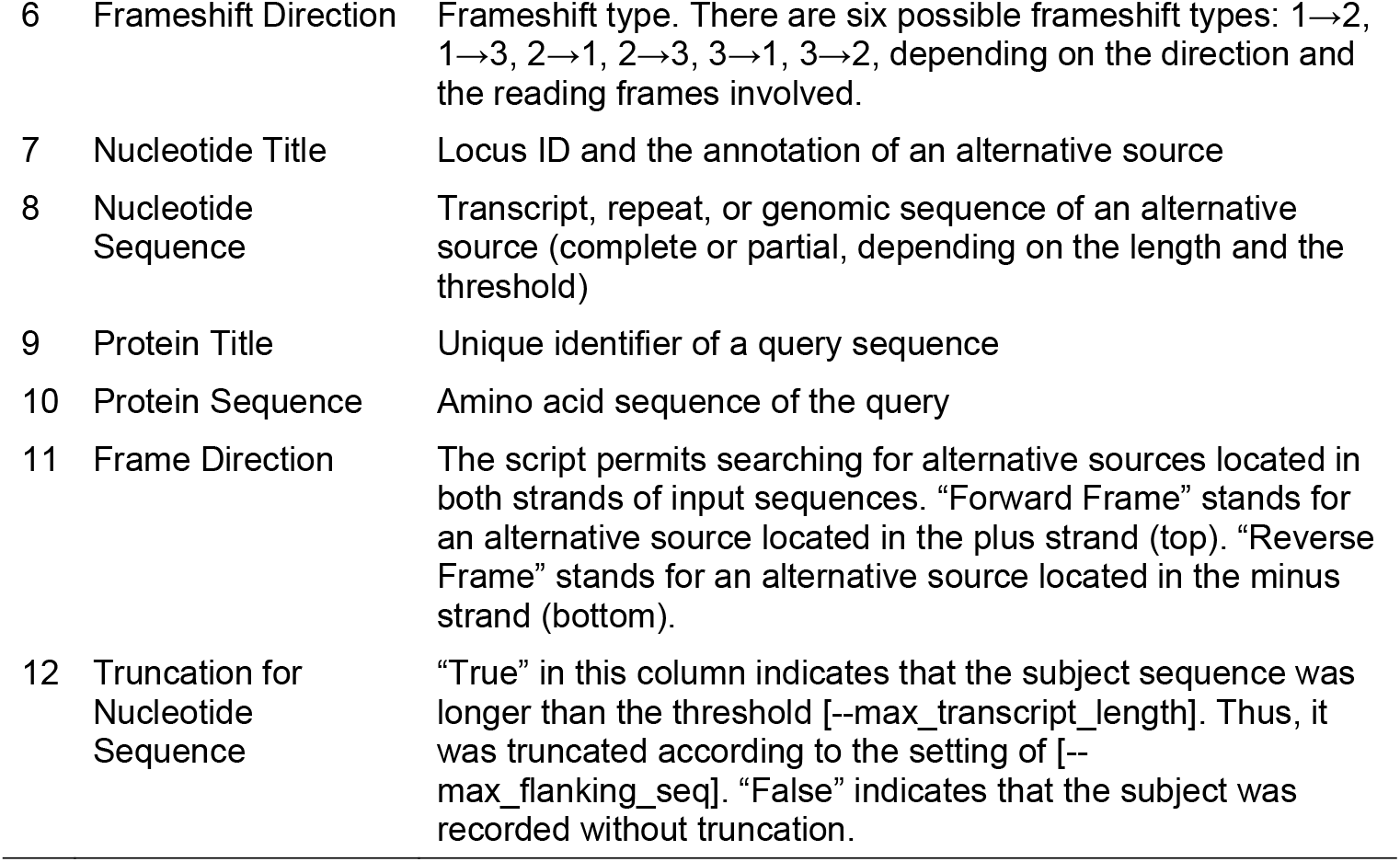
Description of ShiftSCAN outputs.

| Column | Description |
| --- | --- |
| 1 Type | This column indicates whether an alternative source involves a PRF event. "Frameshift" stands for chimeric alternative sources. "Without frameshift" stands for non-chimeric alternative sources. |
| 2 Frameshift Position | This value corresponds to the number of amino acids in a query sequence before the frameshift. |
| 3 Segment 1 | Amino acid sequence before the frameshift |
| 4 Segment 2 | Amino acid sequence after the frameshift |
| 5 Gap | PRF value. Besides the four PRF events considered in our study (-2, -1, +1, and +2), "longer" frameshifts (user-defined, e.g., +10 nucleotides), can also be detected. |

|  |  |  |
| --- | --- | --- |
| 6 | Frameshift Direction | Frameshift type. There are six possible frameshift types: 1→2, 1→3, 2→1, 2→3, 3→1, 3→2, depending on the direction and the reading frames involved. |
| 7 | Nucleotide Title | Locus ID and the annotation of an alternative source |
| 8 | Nucleotide Sequence | Transcript, repeat, or genomic sequence of an alternative source (complete or partial, depending on the length and the threshold) |
| 9 | Protein Title | Unique identifier of a query sequence |
| 10 | Protein Sequence | Amino acid sequence of the query |
| 11 | Frame Direction | The script permits searching for alternative sources located in both strands of input sequences. “Forward Frame” stands for an alternative source located in the plus strand (top). “Reverse Frame” stands for an alternative source located in the minus strand (bottom). |
| 12 | Truncation for Nucleotide Sequence | “True” in this column indicates that the subject sequence was longer than the threshold [--max_transcript_length]. Thus, it was truncated according to the setting of [--max_flanking_seq]. “False” indicates that the subject was recorded without truncation. |

## 3. Results

In *M. truncatula*, we used ShiftSCAN to search for alternative non-chimeric and chimeric sources of 156 MS-supported chimeric peptides. We refer to genetic loci from which these 156 peptides were modeled as primary sources. Three FASTA-formatted lists served as subject databases: (1) the entire genome, (2) the list of all annotated transcripts, and (3) the list of all repeat elements, all downloaded from the current genome release (v. 5.1.9, Pecrix et al., 2018). This analysis revealed that ca. 58% of chimeric peptides (91 out of 156) have unique chimeric sources. Only one chimeric peptide can be produced without any frameshift from genomic DNA. Specifically, it has two non-chimeric alternative loci, both annotated as repeat elements. A separate analysis was based on the TBLASTN search of individual RNA-Seq reads of 50 selected runs downloaded from the Sequence Read Archive (SRA) database (Leinonen et al., 2011; Katz et al., 2022). It showed that four other chimeric peptides can have non-chimeric alternative origin among unusual sequence variants of annotated transcripts. The remaining 60 chimeric peptides have between one and 372 alternative chimeric sources among non-repeat and repeat elements. Careful analysis of expression profiles from the RNA-Seq-based gene expression atlas of *M. truncatula* (Carrere et al., 2021) showed that only 31 chimeric peptides out of 156, which is ca. 20%, have the alternative sources that are at least as likely as their primary sources (Çakır et al., 2024, preprint).

## 4. Discussion

Below, we summarize the advantages of ShiftSCAN exemplified with our own study on PRF in *M. truncatula*. Firstly, the code helped us differentiate between peptides with truly unique chimeric origin and peptides that can potentially be generated via PRF from multiple loci. This is crucial for functional characterization of corresponding loci via loss-of-function approaches. Truly unique loci can be targeted by mutagenesis without the need to generate multiple mutants. Secondly, we found the broad spectrum of alternative chimeric loci, among which there are sequences of non-repeat and repeat type. A group of candidate loci that can potentially produce the same peptide can be targeted by a generic RNA interference construct because alternative loci share considerable sequence similarity. Without the knowledge about alternative loci, the treatment of the wild-type plants with synthetic chimeric peptides would be the only tool available for their functional analysis (Ormancey et al., 2023). Furthermore, with the aid of our software, we identified 70 alternative chimeric sources of repeat type that do not overlap with any other locus at the putative PRF site. Among these loci, ten have expression levels comparable with the expression of non-repeat loci, with log2 TMM values above zero in MtExpress v3 (Carrere et al., 2021). Four of these transcribed repeat elements overlap with no other genes along the whole length of their sequences. Interestingly, the majority of 70 repeat loci mentioned above (41, which is ca. 59%) would require translation from their reverse complements for the production of corresponding chimeric peptides. The information on translated repeat elements is so far very scarce (Davidson et al., 2017; Bourque et al., 2018; Coronado-Zamora and González, 2023), especially in plants (Quadrana et al., 2016). Thus, it deserves dedicated functional studies. Next, our script helped discover chimeric peptides highly conserved among non-repeat and repeat alternative sources, one of which is ultra-conserved among repeat loci. Chimeric peptide CP130 with the primary source annotated as a putative reverse transcriptase, RNA-dependent DNA polymerase, has nine alternative chimeric sources of non-repeat nature and 363 alternative sources among repeat loci. This finding is very intriguing because the reverse transcriptase activity is a characteristic feature of retrotransposons and retroviruses, both known to use PRF for translation (Xiong and Eickbush, 1988; Dinman, 2006; Clark et al., 2007; Riegger and Caliskan, 2022). The software also made it possible to quantify the conservation of PRF sites at the protein level. Namely, we found that, despite limited similarity at the nucleotide level and relatively short length (8-30 aa in our study), MS-supported chimeric peptides with alternative sources exhibit almost no deviation from their primary sources with regard to the PRF value and the frameshift position in the peptide. For example, an MS-validated chimeric peptide CP60 (RCL→GVGARR) was modeled from locus MtrunA17_Chr3g0135761 (primary source) annotated as a putative peptidase. It involves an event with PRF value +2 that shifts from frame 2 to frame 1 (PRF type 2→1). The frameshift position in the peptide is 3 (L→G). This peptide can potentially be produced by exactly the same type of event (PRF value: +2, PRF type: 2→1, position in the peptide: 3) from three other loci annotated as putative peptidases: MtrunA17_Chr1g0158881, MtrunA17_Chr3g0135721, and MtrunA17_Chr8g0362761. Thus, the sequence around the PRF site appears to be conserved in almost all cases of multiple sources. In fact, only four chimeric peptides out of 156 have either PRF value or position (or both) different from their primary sources. For example, an MS-validated chimeric peptide CP37, sequence FLLNLH→QK, was identified as the product of a backward PRF event (−1, 2→1, position 6) from its primary source MtrunA17_Chr2g0299561 annotated as a putative disease resistance protein. Our software showed that it can potentially be produced from a repeat locus MtrunA17_Chr4R0154340 as a peptide FLLN→LHQK (a forward PRF event with parameters +2, 3→2, position 4), but also from another repeat locus, MtrunA17_Chr4R0290860, as a peptide FLLNLHQ→K involving a backward PRF event with parameters -2, 1→2, position 7. Most MS-validated chimeric peptides in our study either have no alternative source or have an alternative source with a PRF site perfectly conserved at the protein level. This is surprising because short chimeric peptides can be reasonably expected to have less unique nature compared to non-chimeric sequences of the same length. For example, 28 chimeric peptides in our dataset contain only one amino acid in the shorter segment before or after a corresponding PRF site. Since we considered four different PRF values in our analysis, the presence of only one amino acid from a different reading frame could theoretically be explained in many different ways, depending on how abundant that amino acid is around the putative PRF site. Contrary to that expectation, our software revealed that 13 of those 28 chimeric peptides have unique sources. It was also surprising to find that seemingly non-unique very short chimeric peptides, e.g., 8 aa-long, can have a single source, as exemplified by an MS-validated chimeric peptide CP146 (ILDTHI→HR). This peptide can be produced exclusively from locus MtrunA17_CPg0493291 by a forward PRF event with parameters +2, 2→1, position 6. Lastly, because the primary purpose of our study was to find evidence for mosaic translation, we searched for transcripts associated with multiple PRF events. Before the application of ShiftSCAN, we found eight such multi-PRF transcripts, all of non-repeat type. Two of those transcripts are likely candidates for mosaic translation. The software enabled the identification of additional seven non-repeat loci and eight repeat loci possibly associated with multiple PRF events. Although this expanded knowledge on multi-PRF loci lowered the probability of mosaic translation in the two best candidate loci, it provided a more realistic view, which is crucial in research on hypothetical processes such as mosaic translation in non-viral systems.

## 5. Conclusions

ShiftSCAN facilitated the generation of biologically relevant information that has not been available before. It also provided evidence that will help focus future functional studies on individual loci or subsets of loci that could potentially serve as templates for PRF. We hope that the evident advantages of this software will motivate researchers to integrate it into their routine analytical pipelines. For example, it would be highly informative to employ this software in the follow-up study on 403 recently discovered chimeric peptides in humans (Ren et al., 2024). We believe the code will be instrumental when the large-scale detection of PRF events in cellular life forms becomes less challenging.

## Abbreviations

altORF: alternative open reading frame
refORF: reference open reading frame
altProt: alternative protein
refProt: reference protein
PRF: programmed ribosomal frameshifting
mRNA: messenger RNA
ncRNA: non-coding RNA
rRNA: ribosomal RNA
tRNA: transfer RNA
aa: amino acids
nt: nucleotides
MS: mass spectrometry

**Umut Çakır:** Conceptualization, Methodology, Software, Validation, Formal analysis, Resources, Data Curation, Writing - Original Draft, Writing - Review & Editing, Visualization, Funding acquisition. **Noujoud Gabed**: Conceptualization, Validation, Formal analysis, Resources, Writing - Review & Editing. **Ali Yurtseven**, Methodology, Software, Formal analysis, Writing - Review & Editing. **Igor S. Kryvoruchko**: Conceptualization, Methodology, Validation, Formal analysis, Resources, Data Curation, Writing - Original Draft, Writing - Review & Editing, Visualization, Supervision, Project administration, Funding acquisition.

## Data availability

The source code for the ShiftSCAN software, along with the documentation and example datasets, is available from the following GitHub repository and PyPI project: https://github.com/umutcakir/shiftscan https://pypi.org/project/shiftscan

## Acknowledgements

This work was supported by the Scientific and Technological Research Council of Turkey (TÜBİTAK) grants and Boğaziçi University standard research grant (BAP-P) to UÇ and IK (TÜBİTAK 1001 120Z514, TÜBİTAK 1002 120Z247, and BAP-P 18841). Computational analysis was conducted using the server of the Turkish National e-Science e-Infrastructure (TRUBA) center. The completion of this study was possible due to the support of IK by United Arab Emirates University and the support of UÇ by the IMPRS-Genome Science PhD program.

